# Do lovebirds ever forget? Long-term memory retention and adaptive forgetting in rosy-faced lovebirds

**DOI:** 10.64898/2026.07.16.739064

**Authors:** Shengyu Wang, Jackson Wen Jie Li, Chen Ma, Emily Shui Kei Poon, Simon Yung Wa Sin

## Abstract

Long-term memory has been extensively documented in many animals across a range of ecological contexts. Far less is understood, however, about how long memories persist after active modification or how conflicting memories are adaptively regulated to enable flexible behaviour. Parrots are widely recognized for their high intelligence and cognitive capability, yet the study of memory has focused on only a handful of large parrot species. Here, using a binary choice symbolic system, we conducted three experiments on 28 rosy-faced lovebirds (*Agapornis roseicollis*) to investigate (1) memory retention of initial associative learning at intervals of two weeks, half a year, and one year; (2) reversal learning, where the rewarded and unrewarded symbols were swapped; and (3) memory retention, following reversal learning at two-week and half-year intervals. We revealed that (a) initial associative learning memory was strikingly persistent, remaining detectable after nearly 600 days; (b) reversal learning performance was comparable to that of large parrot species; and (c) while the post-reversal forgetting curve mirrored the pattern of initial learning, it exhibited a shorter retention duration and a lower retention plateau. These findings suggest that forgetting may serve as an adaptive mechanism that regulates the retrievability of conflicting information, rather than a simple erasure of memory traces. When the environment changes, the retrievability of previously functional but now maladaptive memories is suppressed, but their long-term storage is preserved for potential future use under analogous circumstances. Such dynamic regulations prevent the repetitive overwriting of information in a fluctuating environment, thereby fostering greater behavioural flexibility across dynamic scenarios.

## 1. Introduction

Animals typically store, retain, and subsequently retrieve acquired information under analogous circumstances, rather than readily discarding it (Shettleworth, 2009). Conserving useful information can save substantial time and energy in relearning, thus outstanding memory is adaptive in various ecological contexts—including improved longevity (Welklin et al., 2024) and reproductive success (Shaw et al., 2019), reduced mortality (McLean et al., 2022), kin-recognition (Mateo & Johnston, 2000), remembering novel foraging skills (Borrego & Dowling, 2016; Briefer et al., 2014; Shaw & Harvey, 2020), cultural continuity (Aplin et al., 2015), migration (Merkle et al., 2019), and food caching and retrieving (Kamil & Balda, 1985). Darwin posits that “mental powers”, including memory, evolved to be adaptive to environmental and social challenges, and developed in tandem with physical attributes (Darwin, 1871). For example, migratory garden warblers (*Sylvia borin*) remember a particular feeding site much longer (at least 12 months) than closely related non-migratory Sardinian warblers (*Sylvia melanocephala momus*) do (two weeks) (Mettke-Hofmann & Gwinner, 2003). Similarly, black-capped chickadees (*Poecile atricapilla*) in Alaska, which face harsher and more unpredictable climate, demonstrate more advanced food-storing techniques than their counterparts in Colorado (Pravosudov & Clayton, 2002).

The encoding, storage, and modification of long-term memory represent a highly sophisticated process. Once stored, memories are stabilized through consolidation, a process that occurs after the initial learning experience (Dudai, 2004; McGaugh, 2000). Within minutes to hours of a learning event, synaptic consolidation occurs, a process essential for memory formation that requires the synthesis of new proteins, a fundamental mechanism across nervous systems (Shettleworth, 2009). However, memory storage may not be a one-time, permanent event. Memory retrieval can return a consolidated memory to a labile state, necessitating an active process of re-stabilization to maintain the memory after it is retrieved (Misanin et al., 1968). This dynamic process has been termed reconsolidation, which is an essential module for maintaining long-term memories, allowing for potential updating, strengthening or modification (Dudai, 2006; Tronson & Taylor, 2007). A study by Boccia et al. (2005) demonstrated that exposing mice to a novel learning task immediately, but not three hours, after retrieval of an inhibitory avoidance memory impaired retention performance, supporting the idea that retrieved memories become labile immediately after retrieval, and thus require reconsolidation to be maintained.

While it may appear that natural selection would favour good memory, preserving memories accurately over extended durations can also be costly, considering the massive expenses of maintaining and repairing neural circuits (Dukas, 1999). Clutching to invalid old memories can also lead to maladaptive behaviours when environmental conditions shift. Thus, forgetting obsolete information can be beneficial; from insects to mammals, animals have evolved to forget actively (Cervantes-Sandoval et al., 2016; Epp et al., 2016; Frankland et al., 2013; Hardt et al., 2013; Shuai et al., 2010). As the significance of memory pertaining to specific events wanes, such as condition changes or prolonged absence of similar contexts, such memory will likewise undergo a process of forgetting. Accessibility of memory for a particular item is associated with the frequency, recency, and pattern of its prior exposures (Anderson & Schooler, 1991).

The capacity to flexibly update memories and adjust behaviour is critical for adapting to changing environment in various ecological scenarios (Foley et al., 2008; Grant & Grant, 1989; Reader & Laland, 2003; Sol et al., 2002). It is demonstrated by juvenile cactus finches (*Geospiza conirostris*) during a severe drought; those that successfully learned to forage on novel food sources had a significantly higher survival rate than those who persisted with the foraging behaviours typical of the wet season (Grant & Grant, 1989). Rather than being a generalized attribute, behavioural flexibility is more likely to be exhibited by animals inhabiting complex and unpredictable environments. Flexible animals demonstrate a capacity for rapid behavioural adaptation, drawing on limited experience to respond to subtle contextual or consequential changes (Easton, 2004). A common metric for evaluating behavioural flexibility in psychological and neuroscientific research is reversal learning (Bond et al., 2007; Cauchoix et al., 2017; van Horik & Emery, 2018). This paradigm involves learning to differentiate between a rewarded and a non-rewarded stimulus. After this initial association is firmly established, the reward contingencies are inverted (for instance, the previously rewarded stimulus is no longer rewarded, and vice versa). This shift forces the subject to suppress the original, now-incorrect learning and form a new behavioural response based on the updated rules. Subjects typically struggle immediately after the switch, making numerous errors due to interference of prior learning, i.e., proactive interference (Kane & Engle, 2000). Subsequently, performance gradually recovers and stabilizes over multiple reversal sessions.

Nevertheless, the mechanisms of memory updating and the influence of past experience on present behaviour remain poorly understood due to limited empirical evidence. Evolving theories offer different explanations. Trace decay theory proposes that memories are stored as physical traces in the brain that naturally deteriorate over time, leading to forgetting unless they are actively maintained through rehearsal (Altmann & Gray, 2002). Early interference theory proposed that one memory could prevent the retrieval of another through a process of displacement, wherein new memories essentially overwrite conflicting older ones (Melton & Irwin, 1940). A more nuanced view, known as the retrievability mechanism, suggests that old memories are actively suppressed and regulated, rather than substituted (Kraemer & Golding, 1997). In this learning and forgetting paradigm under changing conditions, the same stimulus can trigger both the obsolete memory (X) and the new memory (Y); thus, adaptive behaviour requires a dual process: encoding new information (Y) while actively suppressing the old one (X). This inhibitory mechanism is vital to prevent behavioural perseveration and facilitate cognitive flexibility.

While sophisticated cognitive abilities were once thought to be exclusive to primates and other mammals, research has revealed that certain bird species, particularly parrots and corvids, possess comparable skills—earning them the nickname “feathered apes” (Lambert et al., 2019). These species demonstrate exceptional capabilities such as innovation, tool use, social learning, and inference (Lambert et al., 2019; Rogers & Kaplan, 2004). A leading hypothesis for the emergence of such complex behaviours in both birds and primates is that they are mediated by an efficient long-term memory, which allows animals to recall specific events and associate them with appropriate behaviours (Chase & Heinemann, 2001; Fagot & Cook, 2006; Rosch, 1978). However, cognitive studies have predominantly been confined to a limited number of large parrot species (Auersperg & von Bayern, 2019). It therefore remains under-explored whether small parrot species, despite their smaller absolute body and brain sizes, exhibit similarly advanced cognitive performance.

Rosy-faced lovebirds (*Agapornis roseicollis*) are highly social, small-sized parrots native to arid regions of southwestern Africa, where seasonal variations lead to substantial fluctuations in food resources (Huynh et al., 2023; Ndithia & Perrin, 2006b). In this challenging environment, they forage socially on a highly selective diet (Ndithia & Perrin, 2006a). According to the cognitive buffer hypothesis, such complex ecological and social pressures are expected to drive the evolution of intelligence (Allman et al., 1993), a premise supported by other frameworks like the Social Intelligence Hypotheses (Humphrey, 1976; Jolly, 1966) and Cultural Intelligence Hypotheses (Whiten & van Schaik, 2007). They are also popular companion pets (Chan et al., 2021), which facilitates a large sample size. These make lovebirds excellent subjects for the study of their long-term memory and reversal learning.

In this study, we investigated the long-term memory of rosy-faced lovebirds, and determined whether the reversed memory was as robust as the original. We first trained the birds on a visual discrimination task and systematically tested the retention of this memory over gaps of two weeks, six months, and one year. Having established a baseline for memory longevity, we then determined if the birds could flexibly update this memory through reversal learning. We also tested the retention of the reversed memory after two weeks and six months, to compare the forgetting curve of the initial memory with that of the newly acquired reversed memory after an identical retention interval. This direct comparison provides a powerful test of the mechanisms underlying memory consolidation and updating, offering new insights into how learned information is maintained and altered over time.

## 2. Materials and Methods

### 2.1 Study species and housing condition

This study was based on 28 adult rosy-faced lovebirds (19 males and 9 females). They were housed separately in wire-mesh cages (length × width × height: 60cm × 40cm × 40cm), each equipped with two perches (60 cm and 25 cm in length), a wooden platform, a swing, a climbing ladder, two chewable wooden blocks, and a variety of naturally sourced enrichments, including branches, straw, pine cones, cuttlefish bone, dried loofah, and dried lotus pods. The light/dark cycle of their room is 8:00 am - 8:00 pm, with additional lights during 9:00 am - 7:00 pm (6500K illumination; LED T5 tube, 7W, SUNSHINE). The ambient temperature is constant at 22-24°C, and humidity is stable at 50-60%. Artificial food pellets (Mazuri Small Bird Maintenance Diet 56A6) and UV-filtered water were available ad libitum with refilling or changing on a daily basis.

### 2.2 Experimental protocol

The experimental apparatus consisted of a single perch (30 cm long) fitted with two stainless-steel cups (5 cm diameter × 5 cm height) at either end (Fig. 1a and 1b). Bird behaviour was filmed from a top-mounted camera (Mi Camera 2K, Xiaomi Communications Co., Ltd.). Subjects had unrestricted access to both cups. Symbols were printed on the white paper lids (7 cm diameter) sealing the cups, and corresponding symbols were displayed on adjacent vertical signs. The two symbols used were five bars (i.e., ‘|||||’, 23 mm long and 3 mm wide, with a 2 mm gap between them) and an empty circle (i.e., ‘O’, external diameter of 23 mm and internal diameter of 17 mm). Hereafter these symbols are referred to as ‘Symbol 5’ and ‘Symbol 0’, respectively. Furthermore, ‘S5+’ and ‘S0+’ represent the respective conditions in which Symbol 5 or Symbol 0 indicates the food reward.

**Figure 1.**
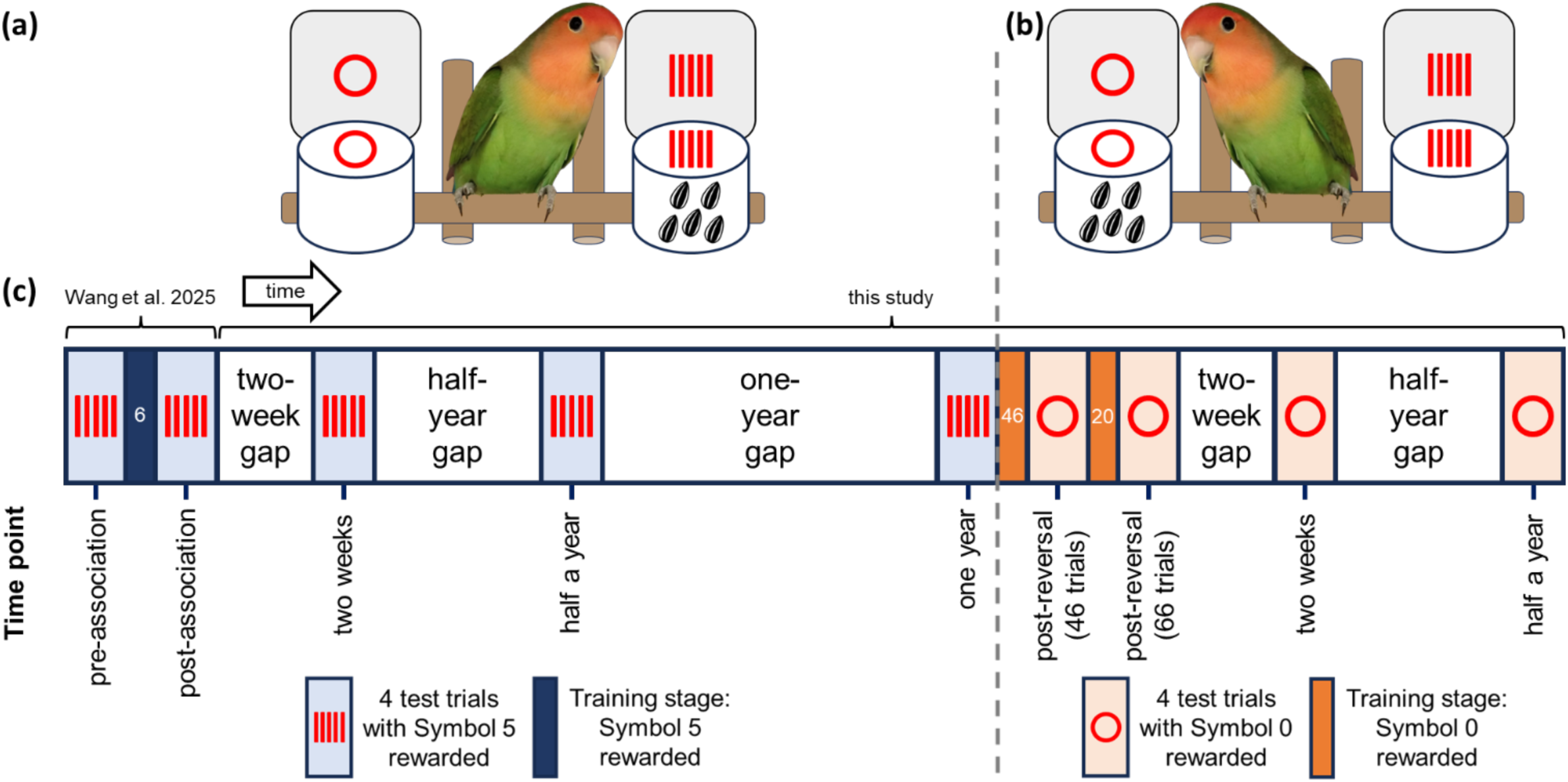
Schematic of the experimental design. (a) Illustration of the trial apparatus for associative learning and subsequent associative learning memory tests. Birds made a choice between two cups, with food reward indicated by Symbol 5. (b) Illustration of the trial apparatus for reversal earning and subsequent reversal learning memory. Birds chose between two cups, with food reward indicated by Symbol 0. (c) Flowchart of the tudy phases, including initial associative learning, associative learning memory, reversal learning, and reversal learning memory tests. Numbers n the training stages indicate the number of trials completed.

The experimental procedures followed those of the previous study (Wang et al., 2025). We limited each bird to one trial per day, with at least one-day interval between trials, to reduce the effects of satiation. To motivate the birds, we closed their feeders 1.5 hours before and 20 minutes after each trial. We then attached both feeder cups to the perch at the same time and allowed the bird 10 minutes to tear the paper lids and consume the food before removing the cups. In the training trials, we removed one-sixth of the paper lid’s area without damaging the symbols, enabling the lovebirds to view the contents to guide their decision while still necessitating effort to tear the lid for access. In the test trials, the cups were completely covered and the birds needed to choose a cup and break the lid before the contents were visible. We considered a choice to be made once the bird had torn the paper off a cup and looked at the content inside. To control for potential side bias (left vs. right) and performance fluctuations at different times of day (morning: 11:00-12:00 vs. afternoon: 16:00-17:00), four test trials were conducted to swap the rewarded symbol position and conduct the trials in the morning and afternoon, for each time point (Table 1; Fig. 1c).

**Table 1.** Experimental timeline and time point definitions. The table details the rewarded stimulus (reward contingency), experimental phases and their purposes, descriptions of each time point with corresponding unique identifiers (time point labels), the test dates of these time points, and the intervals since the previous time point. The time point labels are used consistently throughout the manuscript to denote these specific time points.

| Reward contingency | Phase | Purpose | Time point description | Time point label<sup>a</sup> | Test date | Days since the last time point |
| --- | --- | --- | --- | --- | --- | --- |
| Symbol 5 |  | Establish baseline | Before associative training | Pre-association | 05 Jan - 13 Jan 2023 | NA |
|  | Associative learning | Measure initial learning | After associative training | Post-association | 03 Feb - 10 Feb 2023 | 1 |
|  | Associative learning memory | Measure forgetting | Two weeks after associative training | Two weeks (S5+) | 24 Feb - 04 Mar 2023 | 14 |
|  |  |  | Half a year after the previous test trial | Half a year (S5+) | 12 Sep - 15 Sep 2023 | 192 |
|  |  |  | One year after the previous test trial | One year (S5+) | 14 Sep - 17 Sep 2024 | 365 |
| Symbol 0 | Reversal learning | Measure reversal learning | After 46 reversal training trials | Post-reversal (after 46 trials) | 25 May - 28 May 2025 | 1 |
|  |  |  | After 66 reversal training trials | Post-reversal (after 66 trials) | 02 Jul - 08 Jul 2025 | 1 |
|  | Reversal learning memory | Compare forgetting curves | Two weeks after reversal training | Two weeks (S0+) | 22 Jul - 25 Jul 2025 | 14 |
|  |  |  | Half a year after the previous test trial | Half a year (S0+) | 02 Feb - 05 Feb 2026 | 192 |

<sup>a</sup>S5+ = Symbol 5 indicates food; S0+ = Symbol 0 indicates food.

### 2.3 Initial associative symbol learning (Phase 1)

Associative symbol learning was done in the previous study (Wang et al., 2025): To assess any intrinsic symbol preference, lovebirds were first given four choice trials between cups marked 0 and 5. Birds were then trained to associate 5 with a food reward and 0 with its absence. We did six training trials followed by four test trials to examine the associative learning outcome (Fig. 1c).

2.4 Long-term associative learning memory tests (Phase 2)

Following the initial associative training, we conducted a series of long-term memory retention tests to assess the durability of the learned association and to establish the first forgetting curve. These tests were identical to the previous test sessions but were administered after extended periods without any exposure to the task or stimuli. The first retention test was conducted after a two-week gap from the last test. The second test was administered after a half-year gap from the two-week test. A final test was conducted after a further one-year gap from the half-year test.

### 2.5 Reversal learning (Phase 3)

Following the one-year memory test, the reward contingencies were reversed: Symbol 0 became the rewarded stimulus, and Symbol 5 became non-rewarded. Birds underwent 46 reversal training sessions using the same procedure as the initial associative training, allowing them to inspect the food cup before making a choice. Subsequently, four test trials were conducted to assess their performance. The proportion of first choices for the new rewarded stimulus (Symbol 0) reached approximately 60%. Thus, to reach a level comparable to the post-association test of the initial associative symbol learning (approximately 70%), an additional 20 training sessions were conducted, resulting in 66 reversal training sessions in total. Four test trials were conducted after these reversal training sessions.

### 2.6 Long-term reversal learning memory tests (Phase 4)

To assess long-term memory for the reversed association and to construct a second forgetting curve, a retention test (4 test trials) was administered following a two-week period with no training after the reversal training (66 sessions). Subsequently, a further retention test (4 test trials) was conducted after a six-month interval from the two-week time point.

### 2.7 Data collection

Table 1 provides a summary of each test time point (4 trials), including the corresponding reward symbol, test phase, time point label, test date, and days elapsed since the previous time point. For each trial, we recorded the bird’s first choice (Symbol 0 or 5) and its spatial position (left or right) relative to the subject. Individual performance at each time point was evaluated as the proportion of first choices for the food reward across four trials. General performance at each time point was assessed as the proportion of first choices for the food reward across all trials from all individuals (28 birds × 4 trials = 112 trials per time point). To examine associative learning, associative learning memory, reversal learning, and reversal learning memory, we also computed the average proportion of rewarded first choices for each phase (Phases 1–4; Table 1). We measured the latency to the first choice, defined as the time (in seconds) from the start of the trial to the moment the bird first opened a cup. In addition, we recorded the sex, age, and body weight of each bird (Table S1). Four personality traits—neophobia, exploration, activity, and persistence, obtained from another study (Wang et al., 2026), were included in the analysis to assess their potential influence on task performance (Table S1).

### 2.8 Statistical analysis

We used RStudio (Version 4.2.2) (R Core Team, 2022) for all statistical analyses and data visualization. To assess the effect of symbol position (left vs. right) on the birds’ choice of feeder cup (left vs. right) across all time points, we fitted generalized linear mixed models (GLMMs) with a binomial distribution. These models, implemented using the *lme4* package (Bates et al., 2015), incorporated bird ID and trial ID as random intercepts to account for repeated measures: Model 1) choice position ∼ symbol position + (1|ID) + (1|trial). To compare the proportion of first choice for food rewards across original associative learning and reversed learning performance over identical retention intervals, we conducted Wilcoxon signed-rank tests for three paired time points (Model 2): post-association vs. post-reversal (66 trials); two weeks (S5+) vs. two weeks (S0+); half a year (S5+) vs. half a year (S0+). To evaluate variables associated with individual performance (proportion of first choices for food reward), we ran a generalized linear mixed model (GLMM) with a binomial distribution: Model 3) proportion of first choices for food reward (weighted by total trials) ∼ sex + age + weight + neophobia + exploration + activity + persistence + (1|ID) + (1|time point). Similarly, to evaluate variables associated with latency to the first choice, we ran a linear mixed model (LMM): Model 4) latency to the first choice (log transformed) ∼ sex + age + weight + neophobia + exploration + activity + persistence + (1|ID) + (1|time point). Both Model 3 and Model 4 incorporated bird ID and time point as random effects. Continuous explanatory variables were standardised. In addition, we used Spearman correlation test to investigate the association among associative learning, associative learning memory, reversal learning, and reversal learning memory. Specifically, we examined the correlations between individuals’ proportion of first choices for food reward across these phases (Model 5). All requisite assumptions for the statistical models were examined and satisfied.

#### Ethical note

All procedures were approved by the Committee on the Use of Live Animals in Teaching and Research (CULATR; approval number: 24-202), and under a Department of Health Animal (Control of Experiments) Ordinance Chapter 340 permit ((24-485) in DH/HT&A/8/2/3 Pt.70). All birds were housed in individual cages within a shared aviary, allowing for social contact while preventing aggressive encounters over territory. To promote welfare, we provided environmental enrichment, including toys, chewable objects, and classical music. The cognitive tasks served as additional enrichment, enhancing the birds’ overall well-being. Their health and condition were carefully monitored on a daily basis. We observed no notable behavioural abnormalities linked to food deprivation or isolated caging. Our sample size (n = 28) mirrored that of prior studies using similar designs (Tsang et al., 2025; Wang et al., 2025), which reported significant findings with equivalent numbers. All lovebirds were sourced from local breeders and transferred to our animal facility via transport cages within an hour. After completing this study, the birds were retained for subsequent projects. All protocols adhered fully to the current ASAB/ABS “Guidelines for the ethical treatment of nonhuman animals in behavioural research and teaching” (ASAB Ethical Committee/ABS Animal Care Committee, 2026).

## 3. Results

### 3.1 Long-term retention of initial association

In the previous study (Wang et al., 2025), the birds’ initial proportion of first choices for the rewarded cup was 49.53% before training, which did not differ from chance level (Table 2; Model 1: *p* = 0.885). After six trials of associative symbol training, their average proportion of first choices for the rewarded symbol significantly increased to 71.17% (Table 2; Model 1: *p* < 0.001).

**Table 2.** Results of Model 1^a^ (set of generalized linear mixed models, GLMMs) testing the effect of symbol position (left / right) on feeder cup choice (left / right) in birds.

| Time point label<sup>b</sup> | Proportion of first choices for food reward (%) | $\beta$ | SE | $p$ | SD of random effect | |
| --- | --- | --- | --- | --- | --- | --- |
|  |  |  |  |  | ID | Trial |
| Pre-association | 49.53 | -0.067 | 0.461 | 0.885 | 1.594 | <0.001 |
| Post-association | 71.17 | 2.249 | 0.550 | **<0.001** | 0.961 | 0.000 |
| Two weeks (S5+) | 67.86 | 2.208 | 0.606 | **<0.001** | 1.505 | 0.538 |
| Half a year (S5+) | 59.82 | 1.278 | 0.531 | **0.016** | 1.756 | 0.479 |
| One year (S5+) | 60.71 | 1.152 | 0.468 | **0.014** | 1.094 | <0.001 |
| Post-reversal (46 trials) | 59.82 | 1.058 | 0.463 | **0.022** | 0.789 | <0.001 |
| Post-reversal (66 trials) | 71.43 | 3.127 | 0.760 | **<0.001** | 2.052 | <0.001 |
| Two weeks (S0+) | 56.25 | 0.726 | 0.468 | 0.121 | 1.238 | <0.001 |
| Half a year (S0+) | 57.14 | 0.717 | 0.431 | 0.097 | 1.026 | 0.000 |

<sup>a</sup>Model formula: choice position ~ symbol position + (1|ID) + (1|trial). Significant $p$ -values are in bold.
<sup>b</sup>S5+ = Symbol 5 indicates food; S0+ = Symbol 0 indicates food.

Lovebirds demonstrated significant long-term memory of the initial association, although their performance declined over time, forming a classic forgetting curve (Fig. 2). The mean proportion of first choices for the initially rewarded stimulus fell to 67.86% after a two-week interval, a performance level that remained significantly above chance (Table 2; Model 1: β = 2.208, SE = 0.606, *p* < 0.001). After half a year, performance decreased further, with the proportion of first choices for rewards dropping to 59.82%, which was still significantly above chance (Table 2; Model 1: β = 1.278, SE = 0.531, *p* = 0.016). Performance then appeared to stabilize, with subjects choosing the initially rewarded stimulus on 60.71% of trials even after a one-year gap, which was also significantly above chance (Table 2; Model 1: β = 1.152, SE = 0.468, *p* = 0.014).

**Figure 2.**
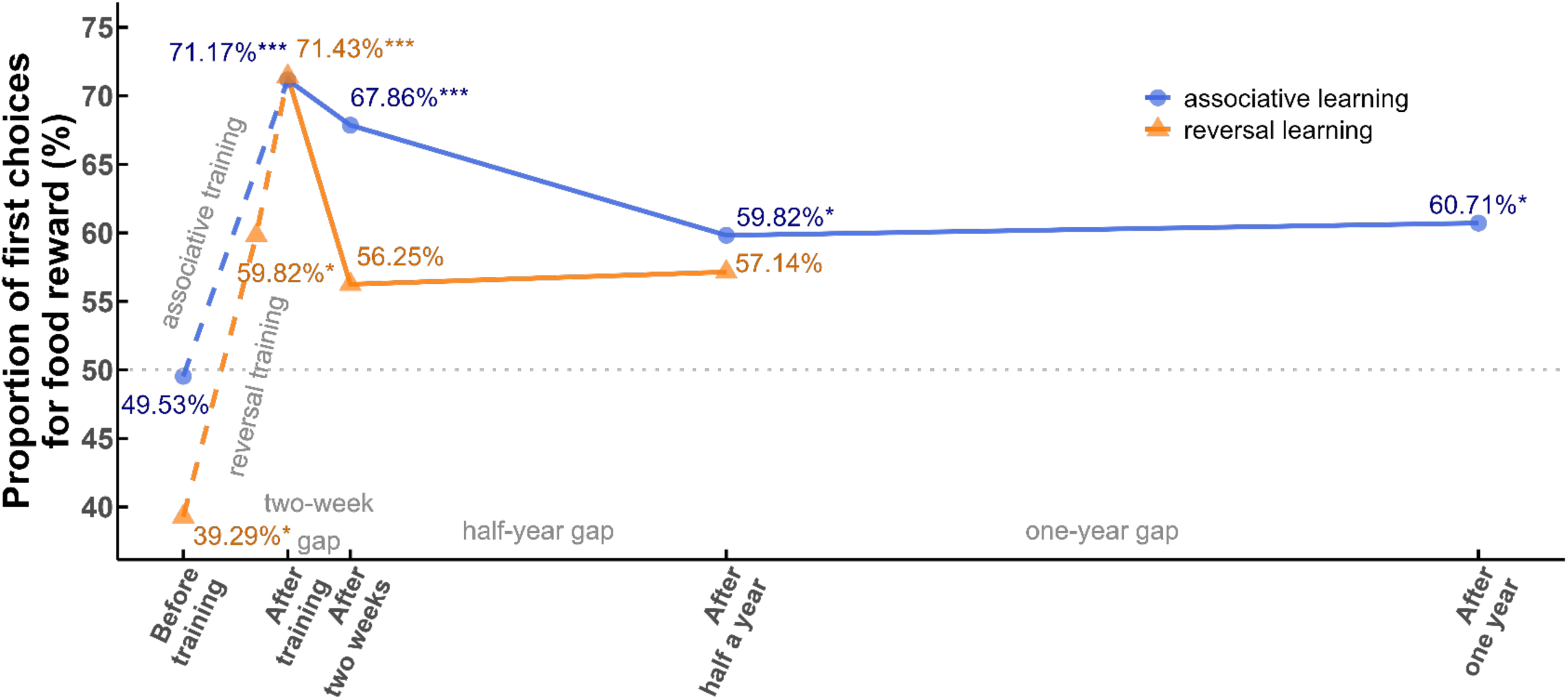
Comparison of forgetting curves for initial and reversed associative memories. The red dashed horizontal line indicates chance erformance (50%). The blue line with circles represents retention of the initial association (Symbol 5 rewarded), tested after a two-week, half-ear, and one-year gap from the previous test trial. The orange line with triangles represents retention of the reversed association (Symbol 0 ewarded), tested after a two-week, and half-year gap from the end of reversal training. Asterisks denote performance significantly above chance * *p* < 0.05, ** *p* < 0.01, *** *p* < 0.001).

### 3.2 Reversal learning

When reward contingencies were reversed, subjects demonstrated the ability to learn the new association (Fig. 2 and Fig. S1). After 46 reversal sessions, the mean proportion of first choices for the newly rewarded stimulus (Symbol 0) reached 59.82%—a level comparable to the performance for the original stimulus (Symbol 5) after one year. This performance was significantly above chance (Table 2; Model 1: β = 1.058, SE = 0.463, *p* = 0.022). Performance was significantly enhanced by an additional 20 sessions of training, increasing the mean proportion of choices for the rewarded stimulus to 71.43% (Table 2; Model 1: β = 3.127, SE = 0.760, *p* < 0.001)—a level comparable to that achieved after initial associative learning training (Table 2, Fig. 2 and Fig. 3a; Model 2: *p* = 0.887).

**Figure 3.**
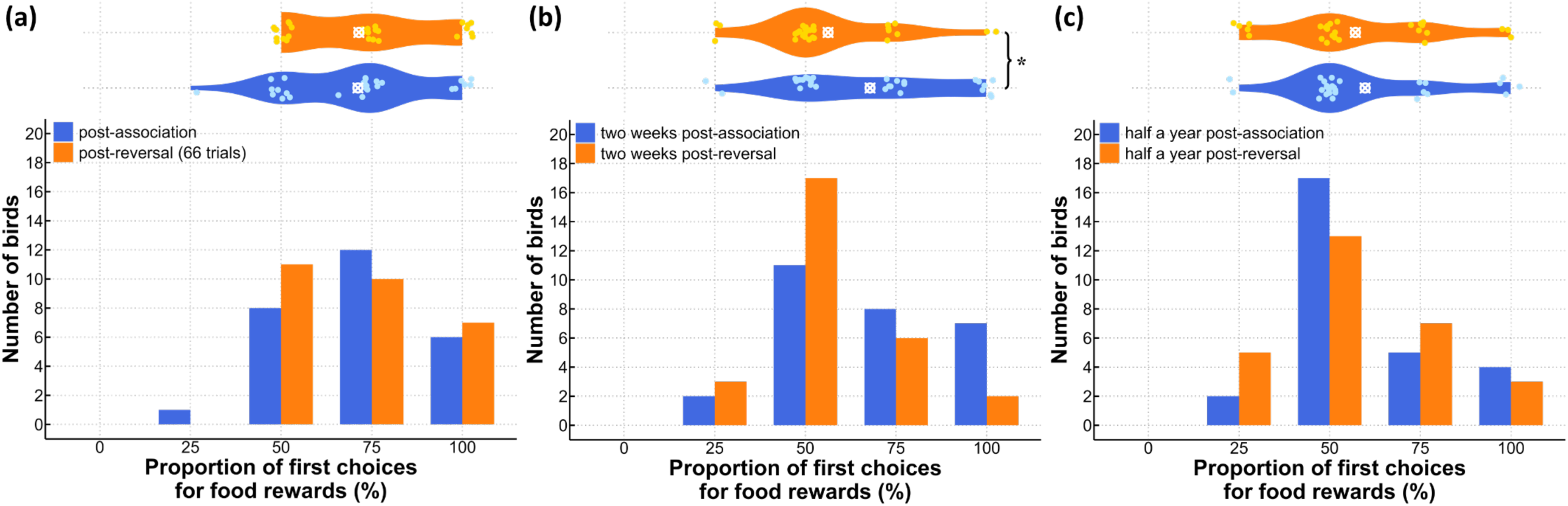
Performance comparison between initial associative learning and reversal learning at identical time point after learning. (a) Comparison f performance after initial associative learning (blue) and after reversal learning (orange). (b) Comparison of memory retention two weeks after he initial associative learning (blue) and the reversal learning (orange). (c) Comparison of memory retention half a year after the initial associative earning (blue) and the reversal learning (orange). Bottom figures are bar plots showing the proportion of first choices for the food reward for each ndividual bird. The superimposed horizontal violin plots show the distribution of these individual proportions. Asterisk denotes a significant ifference in the proportion of first choice for food rewards between two time points (Model 2; * *p* < 0.05).

### 3.3 Memory of the reversed association

Reversal learning memory exhibited a substantially faster rate of forgetting compared to that of initial associative learning memory. After a two-week delay, the proportion of first choices for the newly rewarded stimulus fell to 56.25%, which did not significantly differ from chance levels (Table 2; Model 1: β = 0.726, SE = 0.468, *p* = 0.121). This memory for the reversed association after two weeks was significantly weaker than that for the initial association memory after two weeks (Table 2, Fig. 2 and Fig. 3b; Model 2: *p* = 0.043). After six months, the proportion of first choices for the reversed rewarded stimulus was 57.14%, which was also not significantly different from chance (Table 2; Model 1: β = 0.717, SE = 0.431, *p* = 0.097). This performance was comparable to the retention of the initial association memory after half year (Table 2, Fig. 2 and Fig. 3c; Model 2: *p* = 0.657).

### 3.4 Effects of individual traits on choice proportion and latency

None of the examined variables—sex, age, body weight, neophobia, exploration, activity, or persistence—significantly predicted the proportion of first choices for food reward (Table S2; Model 3). However, among these variables, neophobia had a significant effect on the latency to make the first choice, with more neophobic individuals taking longer to choose (Table S2; Model 4: *β* = 0.223, *p* = 0.032).

### 3.5 Correlations among different cognitive processes

Among all correlations, only associative learning memory and reversal learning memory showed a significant positive correlation (ρ = 0.415; *p* = 0.028). No significant correlations were found between any of the other cognitive processes: associative learning and associative learning memory (ρ = -0.108; *p* = 0.583); associative learning and reversal learning (ρ = -0.242; *p* = 0.215); associative learning and reversal learning memory (ρ = 0.063; *p* = 0.749); associative learning memory and reversal learning (ρ = -0.024; *p* = 0.903); and reversal learning and reversal learning memory (ρ = 0.238; *p* = 0.223) (Fig. S2; Model 5).

## 4. Discussion

Our study demonstrates that while rosy-faced lovebirds possess a capacity for long-term memory, the retention duration of memory is critically dependent on the learning context in which it was acquired (i.e., initial vs. reversed associations). We showed that an initial association is remarkably durable, remaining detectable even after a full year. When this established memory was subsequently updated through reversal learning, the forgetting curve followed a pattern similar to that of the initial association—characterized by a rapid decline before stabilizing. However, the new, reversed association proved to be significantly more vulnerable to forgetting. This asymmetry in memory strength was reflected in three key aspects: compared to the initial association, recall of the reversed association exhibited (a) a substantially faster rate of forgetting over the same time interval (two-week retention: 67.86% vs. 56.25%), (b) shorter overall retention (> 580 days vs. < 14 days), and (c) a relatively lower level of recall during the long-term stable stage (∼ 60% vs. ∼ 57%). This direct comparison reveals that a memory formed through the updating of a prior well-consolidated one is not as robust, offering new insights into the dynamics of memory retrievability across learning scenarios.

Our results reveal a remarkably long-lasting memory in rosy-faced lovebirds for the association between abstract symbols and the presence or absence of food. This associative learning memory persisted for at least one year and a half, with no noticeable decay between the final two time points (i.e. from half-year to one-year). This indicates the formation of a highly stable long-term memory trace. The longevity of this retention is particularly striking given that it was established through minimal associative training of only six trials. Over the memory testing period, the lovebirds’ memory likely underwent processes of synaptic consolidation and reconsolidation (Dudai, 2004, 2006), resulting in a robust engram that showed little degradation over the one-year interval. Such memory duration is comparable to that demonstrated by larger parrot species. For example, the kea (*Nestor notabilis*) can remember a conditioned aversion to a repellent bait for nearly one year (McLean et al., 2022). Orange-winged Amazon parrots (*Amazona amazonica*) demonstrated long-term memory by retaining the learned rules of the Hamilton search task for a period of six months (Cussen & Mench, 2014). Similarly, budgerigars (*Melopsittacus undulatus*), another small parrot, can remember the unique acoustic features of numerous individual contact calls for a period of at least 180 days (Park & Dooling, 1985). Our findings extend this pattern to the domain of visual symbol association in a small parrot, suggesting that the capacity for certain forms of long-term memory may be highly conserved across the parrot family, independent of body and brain size. It is important to note, however, that memory is not a unitary faculty. Various memory systems (e.g., spatial or episodic), shaped by different ecological demands and evolutionary history, may involve distinct cognitive processes, as seen in Clark’s nutcrackers (*Nucifraga columbiana*) (Olson et al., 1995), dark-eyed juncos (*Junco hyemalis*) (Cristol et al., 2003), white-crowned sparrows (*Zonotrichia leucophrys*) (Pravosudov et al., 2006), and Western scrub jays (*Aphelocoma californica*) (Pravosudov et al., 2005). Thus, superior performance on one type of memory task does not necessarily predict equally strong abilities on others.

When placed in a comparative context, the reversal learning performance of rosy-faced lovebirds appears comparable to that of other cognitively renowned birds. Macaws (*Diopsittaca nobilis*) required, on average, approximately 30 trials to learn the initial colour association and 60 for the reversal, while caiques (*Pionites melanocephala*) required around 40 and 75 trials, respectively (van Horik & Emery, 2018). Among corvids, Bond et al. (2007) found that Clark’s nutcrackers, pinyon jays (*Gymnorhinus cyanocephalus*), and Western scrub jays required an average of 5.4, 4.6, and 6.8 sessions (36 trials each), respectively, to reach the learning criterion on their colour reversal. On a spatial task, the mean number of sessions to reach this reversal learning criterion was 2.2, 2.0, and 2.7 for the same species, in the same order. In comparison, research indicates that cats, pigeons, and rats are less proficient at reversal learning, understanding the new contingency slower with inferior flexibility (Durlach & Mackintosh, 1986; MacKintosh & Holgate, 1969; Warren, 1966). While direct numerical comparison across studies requires caution due to methodological differences, the lovebirds’ performance places them firmly within the range of these larger-brained species. This, combined with our previous finding of their long-term memory being comparable to large parrots, strengthens the argument that specific cognitive abilities in small parrots can be on par with those of corvids and large parrots. It suggests that certain cognitive demands, such as tracking changing resource distributions, may select for comparable levels of performance regardless of absolute brain size. Behavioural flexibility, thus, is not a general trait but is often specialized, evolving in response to specific environmental challenges. The demonstrated proficiency in reversal learning by rosy-faced lovebirds, as well as corvids and large parrots, provides strong support for the pliancy hypothesis (Day et al., 1999), which posits that active foraging strategies select for a general cognitive flexibility, or pliancy—the ability to adaptively update stimulus-reward associations. For instance, actively foraging lizards excelled at reversal learning, outperform sit-and-wait foragers (Day et al., 1999). The success of parrots and corvids in reversing a learned discrimination directly exemplifies this type of behavioural flexibility, suggesting their active foraging ecology may have selected for the very cognitive mechanisms that enable rapid adaptation to changing environmental contingencies, as the hypothesis predicts. Their need to track ephemeral resources aligns with the broader pattern, where species in ecologically volatile niches develop more flexible strategies (Davey, 1989; Day et al., 1999), while those in complex physical environments also exhibit greater flexibility (Jones, 2005; Robinson, 1985).

Reversal learning, which involves imprinting contradictory information, did not erase previous memories; instead, they remained encoded in memory repository and were readily recalled in suitable scenarios. In this study, we observed that previous memory not only hindered reversal learning, but also led to accelerated forgetting and reduced retention after reversal (Fig. 2). This indicates a phenomenon of proactive interference, where prior learning interferes with the encoding or retrieval of subsequent new information (Kane & Engle, 2000). To manage this conflict, previous invalid information is temporarily suppressed to prevent its recall when condition changes (i.e., retrieval suppression), while the access to competing memories is adaptively regulated over time according to their recency to optimize behaviour (i.e., retrieval regulation) (Kraemer & Golding, 1997; Sawaki et al., 2012). When retrieval is inhibited by new, competing information, the original memory becomes less accessible but is not erased (Anderson & Huddleston, 2012). Alternatively, if memory updating involved simple erasure or substitution—as trace decay theory proposes—we would expect the two forgetting curves to exhibit similar forgetting rates and comparable retention, given that both began from matched performance levels. Rather than impeding flexibility, proactive interference can be viewed as an adaptive form of forgetting (Kraemer & Golding, 1997), assisting animals manage conflicting memories in dynamic environments. After a long retention interval, a reduced bias toward the recent memory can lead to response uncertainty, allowing an animal to reassess which memory is currently relevant instead of rigidly adhering to a potentially outdated response. This idea is supported by serial reversal learning studies, in which animals improve over successive reversals, reflecting an enhanced ability to adapt to changing contingencies (Bond et al., 2007; van Horik & Emery, 2018).

Our results suggest that the two long-term memory processes we assessed are closely related, but might be mechanistically distinct from associative learning and reversal learning. Although memory for the reversed association was more vulnerable than initial associative learning memory, the two memory processes may share a common underlying neural pathway for encoding and retrieval. This is supported by (a) their similar pattern of change—rapid decrease followed by stabilization—and (b) a positive correlation in individual performance. However, the three cognitive processes examined—initial associative learning, memory retention (associative learning memory and reversal learning memory), and reversal learning—are largely independent in the context of our study (Fig. S2). This lack of correlation points to distinct underlying neural or cognitive mechanisms, challenging the concept of a unitary “learning ability” that underpins different cognitive processes (Galsworthy et al., 2002), and instead supporting a more modular view of cognition (Barrett & Kurzban, 2006). For example, the hippocampus is critical for memory tasks that demand cross-episode integration (Eichenbaum, 2000), whereas the orbitofrontal cortex is implicated in reversal learning (Dalley et al., 2004). Furthermore, the neural basis of memory retention is dynamic, shifting with time from hippocampal dependence to a more distributed cortical network that involves the prefrontal cortex (Frankland & Bontempi, 2005; Kitamura et al., 2017). Empirical evidence suggests that even within a single memory domain, distinct facets such as acquisition speed, accuracy, and retention can be dissociated. For instance, in a radial-arm maze task, Clark’s nutcrackers and pinyon jays learned faster and achieved higher initial accuracy than scrub jays (*Aphelocoma coerulescens*) and Mexican jays (*A. ultramarina*) (Kamil et al., 1994). However, this advantage disappeared after a 300-minute retention interval. These findings indicate that specific processes are dissociable, meaning that initial accuracy, retention, and reversal learning capacity may involve distinct brain regions and evolve independently. Alternatively, the constrained number of trials, which was implemented to limit memory reconsolidation, may have reduced our sensitivity to detect smaller correlations that are typical of complex cognitive traits. Thus, these findings are best interpreted as an absence of large effects rather than conclusive evidence for complete independence.

The enduring yet flexible memory system, which balances robust retention of predictive cues with the ability to adapt to shifting profitability, is ecologically significant given lovebird’s natural history and foraging demands. As highly social foragers in semi-arid environments, where patchy resources fluctuate substantially and periodically (Ndithia & Perrin, 2006a, 2006b), remembering the visual cues associated with profitable feeding sites over extended periods, including seasonal cycles, would be highly adaptive for efficiently relocating ephemeral food sources in their habitat. In this volatile niche, the ability to quickly abandon a now-unproductive foraging site or food type and switch to a newly profitable one would confer a significant adaptive advantage. Moreover, their challenges within a complex social system (Ndithia et al., 2007; Perrin & Laubscher, 2012) require flexible adaptation to dynamic interindividual relationships. Thus, species in permanent social groups, which demand constant behavioural adjustment, tend to show heightened flexibility (Bhumstein & Armitage, 1998; Easton, 2004; Shultz & Dunbar, 2006).

### Conclusion

Our study reveals a critical asymmetry in memory stability: while lovebirds form a durable long-term memory for an initial task, a subsequently learned reversal is significantly more vulnerable to forgetting. These findings suggest that the robustness of a memory is shaped not only by how well it is learned, but also by the regulation between conflicting information. Forgetting, in this context, is not a process of deletion but a temporary inhibition of information retrievability. This work provides key insights into how memory systems are adaptively regulated over time—prioritizing information based on its frequency and recency—to support optimal decision-making, particularly in dynamic environments.

## Data availability

Data and code are available online as supplementary materials.

## Acknowledgements

We would like to express our gratitude to the CCMR staff Dorman T.M. Foo, Mei Ying Wu, and Tung Yau Au for taking care of our lovebirds. We thank Kevin Ching Hei Lo, Maggie Heap Hui Li, Vincy Wing Chi Ng, Christy Yuen Ching Hung, Wynne Wan Hei Ting, Rachel Yung Chan, Swing Lam and Abby Xin Jiang for assistance during the experiments. We are grateful to Maggie Heap Hui Li for her generous contribution of a photograph of her delightful lovebird, TungTung, which was used as the basis for Figure 1a. We thank Charis May Ngor Chan for technical support. The High-Performance Computing (HPC) service provided by the Information Technology Services at the University of Hong Kong supported our data analysis. This work was supported by a Start-up grant from the University of Hong Kong to S.Y.W.S.

## Supplementary Materials

**Table S1.**
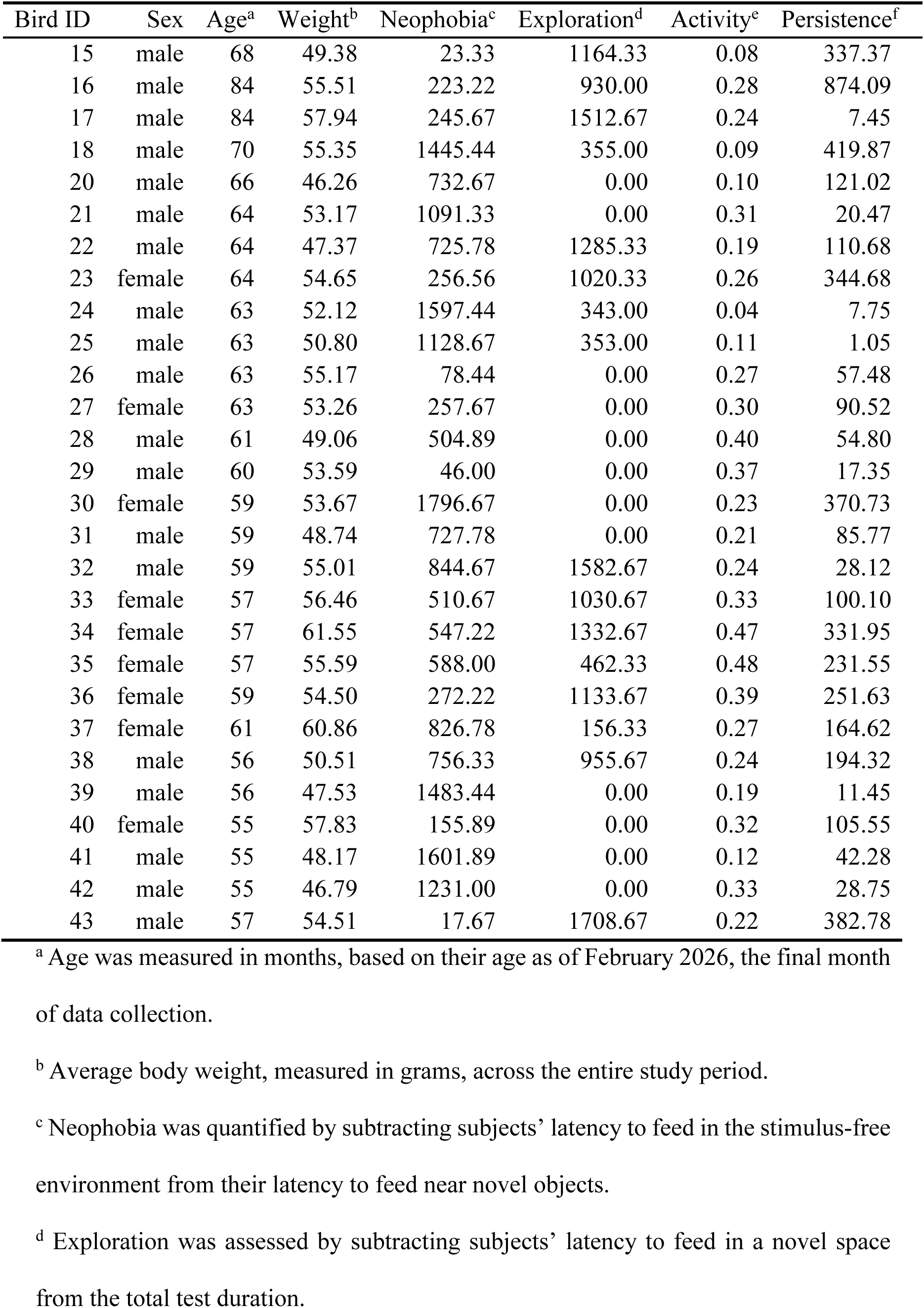
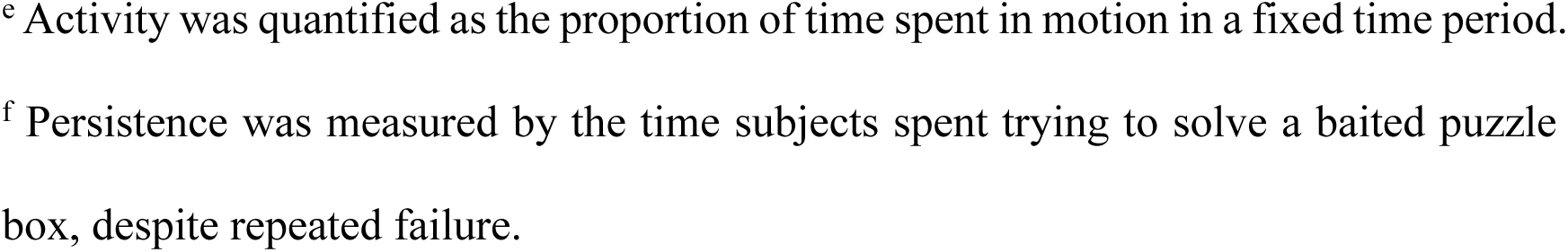
Individual traits.

| Bird ID | Sex | Age<sup>a</sup> | Weight<sup>b</sup> | Neophobia<sup>c</sup> | Exploration<sup>d</sup> | Activity<sup>e</sup> | Persistence<sup>f</sup> |
| --- | --- | --- | --- | --- | --- | --- | --- |
| 15 | male | 68 | 49.38 | 23.33 | 1164.33 | 0.08 | 337.37 |
| 16 | male | 84 | 55.51 | 223.22 | 930.00 | 0.28 | 874.09 |
| 17 | male | 84 | 57.94 | 245.67 | 1512.67 | 0.24 | 7.45 |
| 18 | male | 70 | 55.35 | 1445.44 | 355.00 | 0.09 | 419.87 |
| 20 | male | 66 | 46.26 | 732.67 | 0.00 | 0.10 | 121.02 |
| 21 | male | 64 | 53.17 | 1091.33 | 0.00 | 0.31 | 20.47 |
| 22 | male | 64 | 47.37 | 725.78 | 1285.33 | 0.19 | 110.68 |
| 23 | female | 64 | 54.65 | 256.56 | 1020.33 | 0.26 | 344.68 |
| 24 | male | 63 | 52.12 | 1597.44 | 343.00 | 0.04 | 7.75 |
| 25 | male | 63 | 50.80 | 1128.67 | 353.00 | 0.11 | 1.05 |
| 26 | male | 63 | 55.17 | 78.44 | 0.00 | 0.27 | 57.48 |
| 27 | female | 63 | 53.26 | 257.67 | 0.00 | 0.30 | 90.52 |
| 28 | male | 61 | 49.06 | 504.89 | 0.00 | 0.40 | 54.80 |
| 29 | male | 60 | 53.59 | 46.00 | 0.00 | 0.37 | 17.35 |
| 30 | female | 59 | 53.67 | 1796.67 | 0.00 | 0.23 | 370.73 |
| 31 | male | 59 | 48.74 | 727.78 | 0.00 | 0.21 | 85.77 |
| 32 | male | 59 | 55.01 | 844.67 | 1582.67 | 0.24 | 28.12 |
| 33 | female | 57 | 56.46 | 510.67 | 1030.67 | 0.33 | 100.10 |
| 34 | female | 57 | 61.55 | 547.22 | 1332.67 | 0.47 | 331.95 |
| 35 | female | 57 | 55.59 | 588.00 | 462.33 | 0.48 | 231.55 |
| 36 | female | 59 | 54.50 | 272.22 | 1133.67 | 0.39 | 251.63 |
| 37 | female | 61 | 60.86 | 826.78 | 156.33 | 0.27 | 164.62 |
| 38 | male | 56 | 50.51 | 756.33 | 955.67 | 0.24 | 194.32 |
| 39 | male | 56 | 47.53 | 1483.44 | 0.00 | 0.19 | 11.45 |
| 40 | female | 55 | 57.83 | 155.89 | 0.00 | 0.32 | 105.55 |
| 41 | male | 55 | 48.17 | 1601.89 | 0.00 | 0.12 | 42.28 |
| 42 | male | 55 | 46.79 | 1231.00 | 0.00 | 0.33 | 28.75 |
| 43 | male | 57 | 54.51 | 17.67 | 1708.67 | 0.22 | 382.78 |

<sup>a</sup> Age was measured in months, based on their age as of February 2026, the final month of data collection.
<sup>b</sup> Average body weight, measured in grams, across the entire study period.
<sup>c</sup> Neophobia was quantified by subtracting subjects' latency to feed in the stimulus-free environment from their latency to feed near novel objects.
<sup>d</sup> Exploration was assessed by subtracting subjects' latency to feed in a novel space from the total test duration.
<sup>e</sup> Activity was quantified as the proportion of time spent in motion in a fixed time period.
<sup>f</sup> Persistence was measured by the time subjects spent trying to solve a baited puzzle
box, despite repeated failure.

**Table S2.** Results of LMMs examining predictors of the proportion of first choices for food reward (Model 3) and the latency to make the first choice (Model 4).

| Explanatory variables | Explanatory variables | $\beta$ | SE | z/t | $p$ | SD of random effects | |
| --- | --- | --- | --- | --- | --- | --- | --- |
|  |  |  |  |  |  | bird ID | time point |
| Proportion of first choices for food reward (Model 3) | Sex (male) | 0.136 | 0.210 | 0.648 | 0.517 | <0.001 | 0.223 |
|  | Age | 0.124 | 0.093 | 1.335 | 0.182 |  |  |
|  | Weight | 0.048 | 0.098 | 0.491 | 0.623 |  |  |
|  | Neophobia | 0.106 | 0.083 | 1.269 | 0.204 |  |  |
|  | Exploration | -0.041 | 0.078 | -0.523 | 0.601 |  |  |
|  | Activity | -0.049 | 0.091 | -0.539 | 0.590 |  |  |
|  | Persistence | -0.083 | 0.081 | -1.020 | 0.308 |  |  |
| Latency to the first choice (Model 4) | Sex (male) | 0.038 | 0.246 | 0.153 | 0.880 | 0.366 | 0.274 |
|  | Age | -0.136 | 0.105 | -1.293 | 0.211 |  |  |
|  | Weight | 0.079 | 0.114 | 0.691 | 0.498 |  |  |
|  | Neophobia | 0.223 | 0.097 | 2.301 | **0.032** |  |  |
|  | Exploration | 0.004 | 0.091 | 0.043 | 0.966 |  |  |
|  | Activity | -0.006 | 0.106 | -0.052 | 0.959 |  |  |
|  | Persistence | -0.076 | 0.092 | -0.821 | 0.421 |  |  |

Significant $p$ values are in bold.

**Figure S1.**
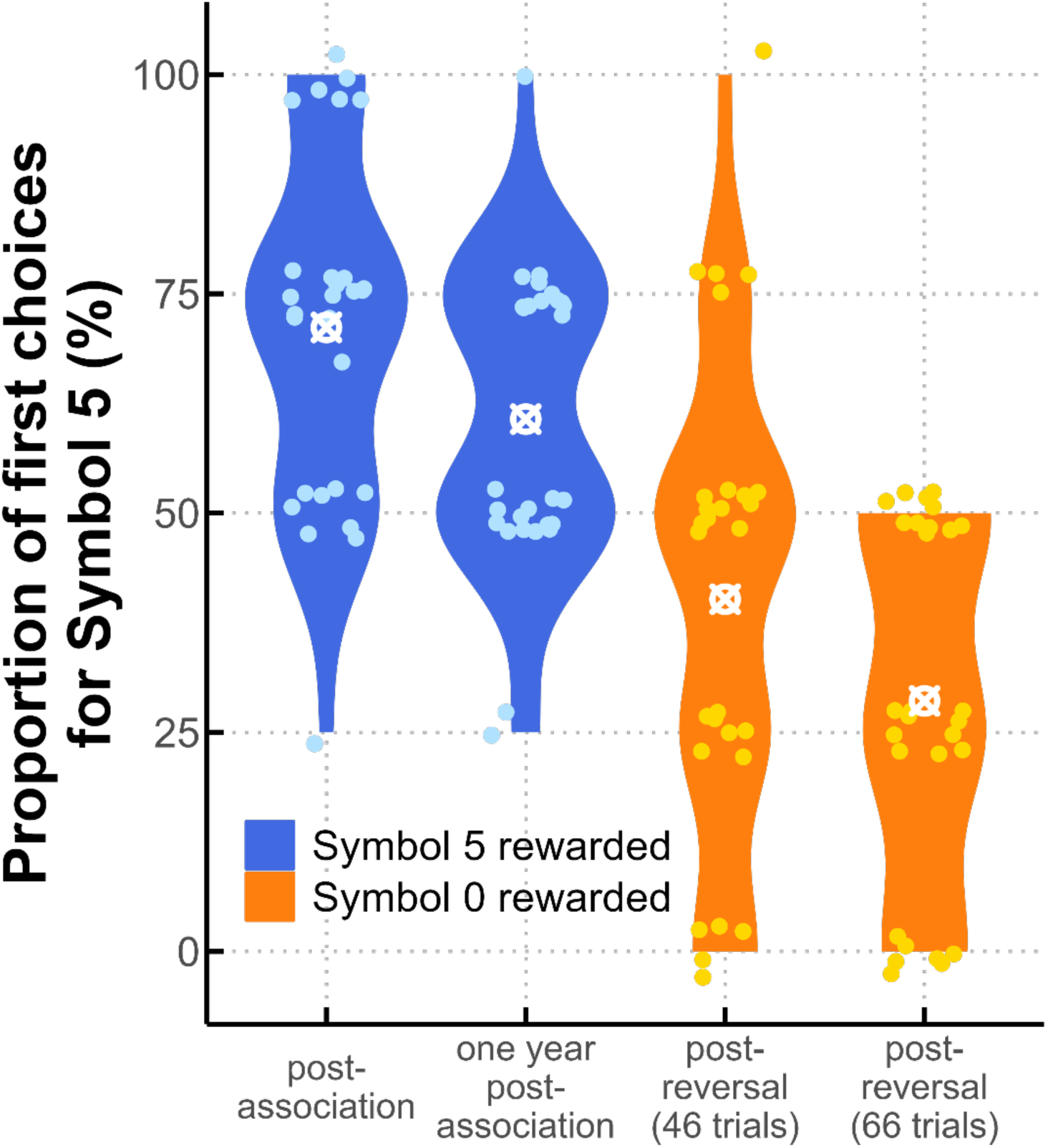
Performance shift across reversal learning. Violin plots show the distribution of the individual proportion of first choices for Symbol 5 across four key time points. Circled crosses indicate the mean proportion for each time point. Blue violins represent time points where Symbol 5 was rewarded; orange violins represent time points where Symbol 0 was rewarded. The proportion for Symbol 5 decreased progressively through reversal training.

**Figure S2.**
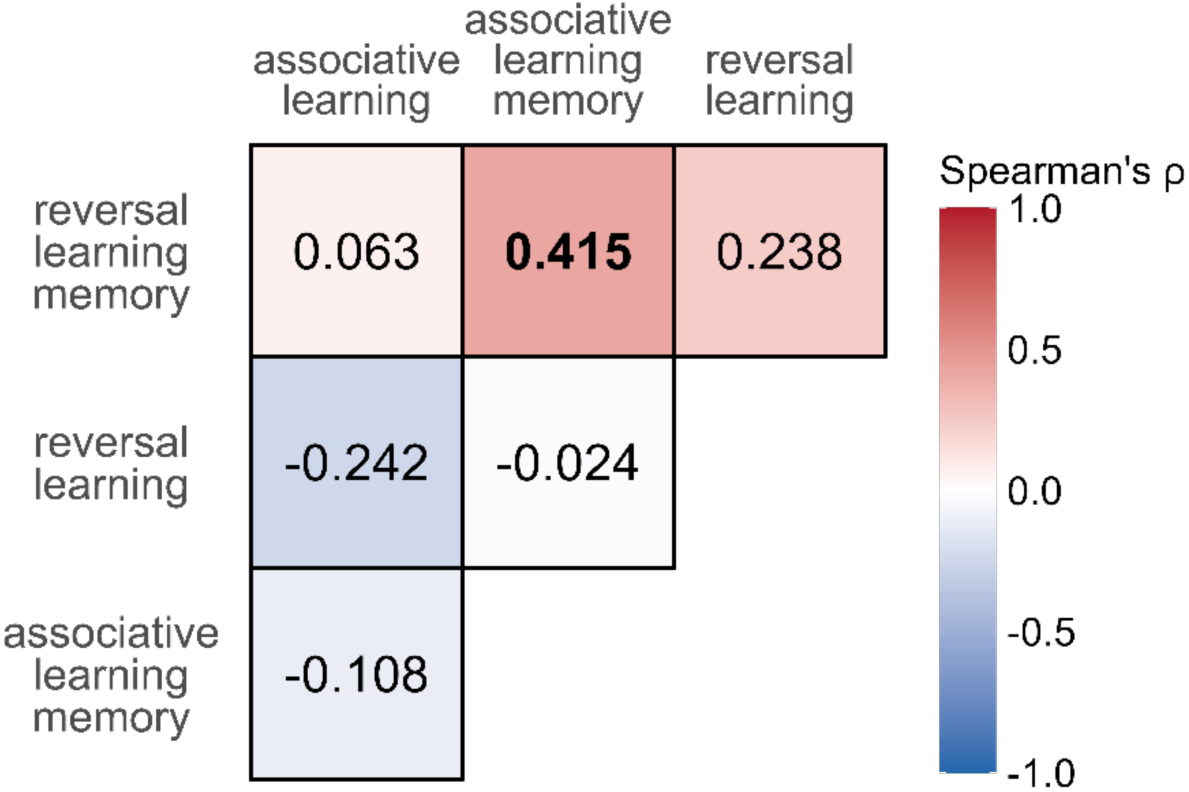
Correlations among cognitive performance measures. Spearman correlation coefficients (ρ) are displayed for each pairwise comparison. Positive correlations are indicated in red, and negative correlations in blue, with colour intensity reflecting the strength of the association (see scale). Significant correlations (*p* < 0.05) are highlighted in bold.

